# Cdt1 variants reveal unanticipated aspects of interactions with Cyclin/CDK and MCM important for normal genome replication

**DOI:** 10.1101/295212

**Authors:** Pedro N. Pozo, Jacob P. Matson, Yasemin Cole, Katarzyna M. Kedziora, Gavin D. Grant, Brenda Temple, Jeanette Gowen Cook

## Abstract

The earliest step in DNA replication is origin licensing which is the DNA loading of MCM helicase complexes. The Cdt1 protein is essential for MCM loading during G1 phase of the cell cycle, yet the mechanism of Cdt1 function is still incompletely understood. We examined a collection of rare Cdt1 variants that cause a form of primordial dwarfism (Meier-Gorlin syndrome) plus one hypomorphic *Drosophila* allele to shed light on Cdt1 function. Three hypomorphic variants load MCM less efficiently than WT Cdt1, and their lower activity correlates with impaired MCM binding. A structural homology model of the human Cdt1-MCM complex position the altered Cdt1 residues at two distinct interfaces rather than the previously described single MCM interaction domain. Surprisingly, one dwarfism allele (*Cdt1-A66T*) is more active than WT Cdt1. This hypermorphic variant binds both Cyclin A and SCF^Skp2^ poorly relative to WT Cdt1. Detailed quantitative live cell imaging analysis demonstrated no change in stability of this variant however. Instead, we propose that Cyclin A/CDK inhibits Cdt1 licensing function independently of the creation of the SCF^Skp2^ phosphodegron. Together, these findings identify key Cdt1 interactions required for both efficient origin licensing and tight Cdt1 regulation to ensure normal cell proliferation and genome stability.

## INTRODUCTION

DNA replication must be tightly regulated to ensure normal cell proliferation throughout development. DNA damage arising from errors in DNA replication can lead to oncogenic transformation, developmental disorders, and aging (Arentson et al. 2002; Blow and Gillespie 2008; Yekezare et al. 2013). The first essential DNA replication step is DNA helicase loading which occurs in G1 phase of the cell cycle through the nucleation of several protein components at presumptive replication origins. This process is known as “origin licensing.” DNA helicase loading renders origins competent for DNA replication in the subsequent S phase. Unscheduled origin licensing after G1 can lead to DNA re-replication, DNA damage, cell death, and genome instability (Vaziri et al. 2003; Melixetian et al. 2004; Li and Jin 2010). For this reason, origin licensing is tightly restricted to G1 to ensure “once, and only once” genome duplication each cell cycle (Cook 2009; Truong and Wu 2011). On the other hand, insufficient licensing increases the probability of incomplete replication, another source of genome instability and proliferation failure (Shreeram et al. 2002; Machida et al. 2005; Nevis et al. 2009).

The Cdt1 (Cdc10-dependent transcript 1) protein is essential for origin licensing in eukaryotic cells. In coordination with ORC (Origin Recognition Complex) and Cdc6 (Cell division cycle 6), Cdt1 recruits and participates in loading the core of the replicative helicase MCM_2-7_ (Minichromosome Maintenance) at presumptive origins. Human Cdt1 licensing activity is restricted to G1 through combinations of transcriptional control, phosphorylation, ubiquitin-mediated degradation, and binding to a specialized inhibitor protein, Geminin (Pozo and Cook 2016). Unlike the ORC, Cdc6, and MCM ATPases, Cdt1 is not an enzyme. Moreover, the Cdt1 primary sequence is not as highly conserved among eukaryotic species as the other licensing factors, and the regulation of human Cdt1 is complex (Fujita 2006). This complexity presumably arose because loss of proper human Cdt1 control is particularly genotoxic (Arentson et al. 2002; Liontos et al. 2007). Extensive biochemical assays of reconstituted yeast origin licensing reactions have demonstrated that Cdt1 is absolutely required for MCM loading, but the precise role of Cdt1 remains relatively mysterious.

We sought additional insight into Cdt1 function by analyzing the consequences of naturally occurring Cdt1 missense alleles. The first partial-loss-of function variant in a metazoan Cdt1 orthologue was described in *Drosophila melanogaster* (Whittaker et al. 2000). The orthologous vertebrate variants had low activity *in vitro* (De Marco et al. 2009; You et al. 2016), but the specific reason for the weak origin licensing activity was not determined. Importantly, several studies have found that inherited mutations in human origin licensing factors, including Cdt1, can result in developmental disorders (Bicknell et al. 2011a; Bicknell et al. 2011b; Burrage et al. 2015). Cdt1 mutations are one cause of a form of primordial dwarfism called Meier-Gorlin Syndrome. Patients are extraordinarily short with microcephaly, focal hypoplasias, and some characteristic facial features and tissue-specific phenotypes (Bicknell et al. 2011a; de Munnik et al. 2012); these phenotypes are consistent with cell proliferation defects. Indeed, primary fibroblasts from Meier-Gorlin Syndrome patients proliferate slowly in culture (Bicknell et al. 2011b). We hypothesized that each of these mutations perturbs at least one aspect of Cdt1 regulation or function. Our analyses of these alleles identifies a previously unappreciated MCM binding site and separately uncovers new features of Cyclin A-dependent Cdt1 control to prevent genotoxic re-replication.

## RESULTS

### Comparative functional analysis of Cdt1 variants by re-replication induction

Bicknell et al. reported 8 *Cdt1* alleles in Meier-Gorlin Syndrome patients (Bicknell et al. 2011a); we marked the positions of the amino acids affected by all missense alleles and one of the three nonsense alleles in Figure 1A. All of the dwarfism patient genotypes were compound heterozygotes, and the most common combinations were a missense allele plus a nonsense allele predicted to encode a truncated Cdt1 protein. We included all missense mutations in our study. In addition, we included *Cdt1-Y520X* because it encodes the longest of the predicted truncations, and we reasoned that if *Cdt1-520X* is null for function then the shorter truncations are also null. We added *Cdt1-R210C*, a variant first discovered as a *D. melanogaster* partial loss-of-function mutant (Whittaker et al. 2000). This analogous vertebrate variant has reduced origin licensing activity *in vitro* (De Marco et al. 2009; You et al. 2016), but the molecular mechanism of reduced activity is not known. We introduced each of these mutations into a vector encoding full length Cdt1 under control of a doxycycline-inducible promoter with a C-terminal extension that includes polyhistidine and HA epitope tags. We then generated derivatives of the U2OS osteosarcoma cell line by recombination of each expression construct into a chromosomal FRT site using Flp recombinase; the parent cell line constitutively expresses the Tet repressor. Using this experimental approach, we achieved dose-responsive inducible ectopic Cdt1 expression (Fig. 1B).

**Figure 1.**
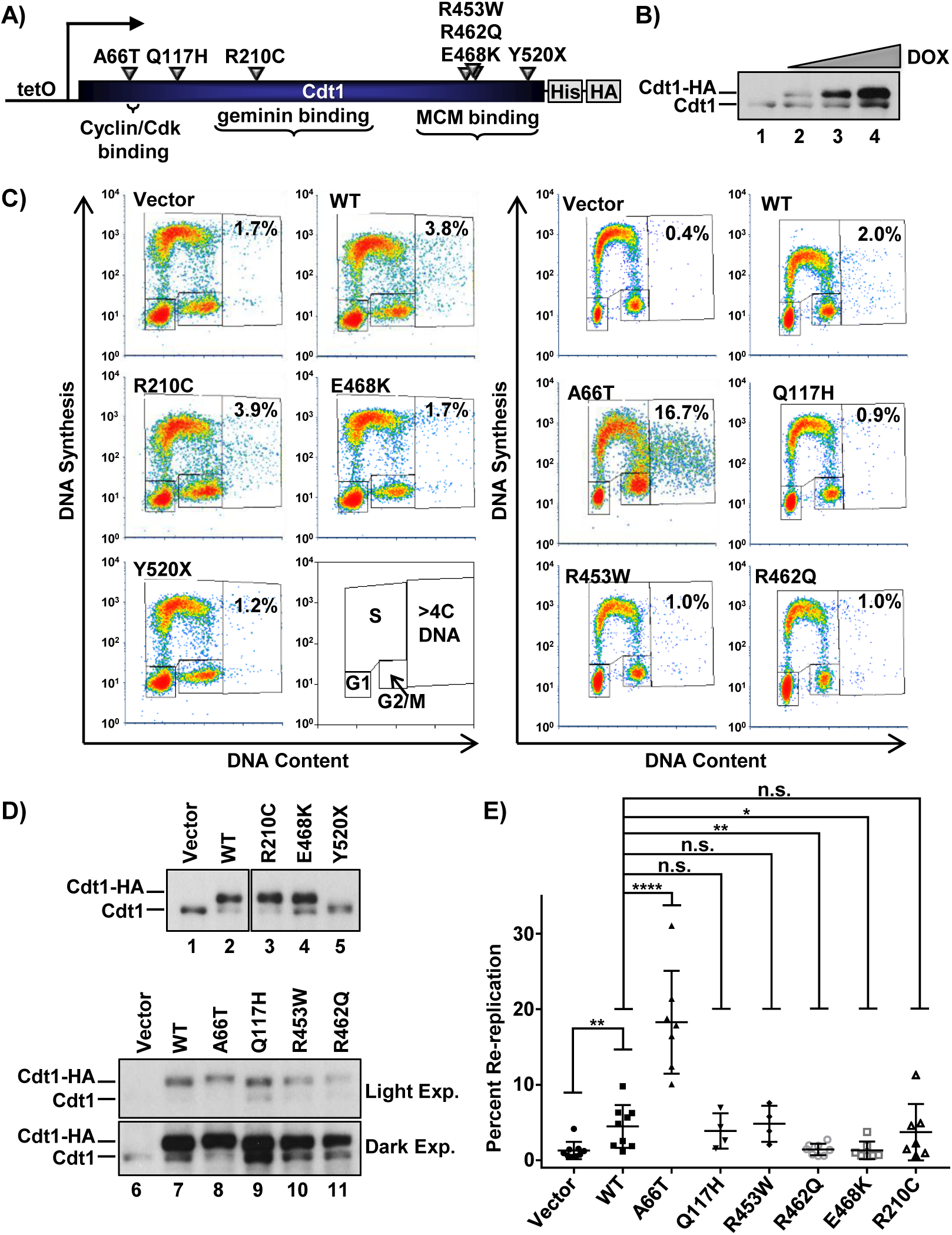
Functional analysis of Cdt1 variants by re-replication induction. *(A)* Illustration of the relative location and amino acid substitution of the alleles chosen in this study; polyhistidine and HA epitope tags and relevant binding domains are also marked. *(B)* Immunoblot of inducible expression of stably-integrated HA-tagged WT Cdt1 in U2OS cells. Cells were grown in 0 µg/mL, 0.05 µg/mL, 0.1 µg/mL, and 1 µg/mL doxycycline (dox), respectively. *(C)* Analytical flow cytometry profiles of U2OS cells expressing either vector, ectopic HA-tagged WT Cdt1, or HA-tagged Cdt1 variants. Cells were treated with 1 µg/mL dox for 72 hrs. and pulse labeled with EdU for 30 minutes prior to harvesting. An illustration of the gating scheme is also shown; “>4C DNA” are cells that have undergone DNA re-replication. *(D)* Immunoblots of Cdt1 expression from *C*. Light Exp. – light exposure; Dark Exp. – dark exposure; spliced images are from the same immunoblot and same exposure *(E)* The percentage of cells with >4C DNA content in at least 4 biological replicates. Bars represent mean and standard deviation. **** = p value < 0.0001; ** = p value <0.005; * = p value <0.05; n.s. = not significantly different.

We first examined the effects of overexpressing each Cdt1 variant by high-dose doxycycline (dox) treatment. Cdt1 overexpression can induce DNA re-replication detectable as a population of cells with greater than the normal G2 phase DNA content (i.e. >4C) (Vaziri et al. 2003). We measured re-replication by analytical flow cytometric analysis after overproducing Cdt1-WT (wild type) or Cdt1 variants in asynchronously proliferating cultures for 72 hrs (Fig. 1C, D). We scored the percent of cells with >4C DNA content in multiple independent experiments (Fig. 1E). Of note, Cdt1 overexpression to this degree had modest effects on the cell cycle distributions among G1, S, and G2/M phases (Supplemental Fig. S1). The more extensive the re-replication, the greater the down-regulation of endogenous Cdt1 which we had previously linked to Cul4-dependent Cdt1 degradation (for example Fig. 1D, endogenous Cdt1 in lanes 7, 8) (Hall et al. 2008).

After multiple independent tests, we noted that Q117H, R210C, and R453W had re-replication-inducing activity similar to WT, whereas R462Q and E468K were less active by this metric (Fig. 1E). Y520X failed to accumulate to high levels at any doxycycline concentration (Fig. 1D, lane 5 and data not shown) which may indicate impaired protein folding and by extension, that all truncation alleles are likely null for Cdt1 biological activity. Given the proliferation defect associated with Meier-Gorlin Syndrome, we anticipated that most alleles encode partial loss-of-function variants like R462Q and E468K. Surprisingly however, the A66T dwarfism variant consistently induced nearly four-fold more re-replication than Cdt1-WT did (Fig. 1C, E), even when produced to similar levels (Fig. 1D), indicating that it is a gain-of-function allele. We focused our subsequent analyses on the subset of mutations with detectable effects on Cdt1 activity *in viv*o, i.e. A66T, R462Q, E468K (Fig. 1C, E), and R210C (Whittaker et al. 2000; De Marco et al. 2009; You et al. 2016).

Given that DNA re-replication is associated with DNA damage and genomic stress, we assessed Cdt1-overproducing cells for activation of the DNA-damage response. We analyzed the activating Chk1 phosphorylation at S345 as a marker of replication stress and DNA damage 48 hrs after initiating Cdt1 overproduction (Fig. 2A, B). As expected (Vaziri et al. 2003; Davidson et al. 2006; Hall et al. 2008), Cdt1-WT overexpression induced Chk1 phosphorylation that correlated with the degree of re-replication induced by the different Cdt1 variants (Fig. 2B). Of particular note, Cdt1-A66T induced significantly more re-replication and Chk1 activation than Cdt1-WT or Cdt1-R210C did, whereas Cdt1-R462Q and Cdt1-E468K induced significantly less re-replication and Chk1 activation than Cdt1-WT did (Fig. 2A, B).

**Figure 2.**
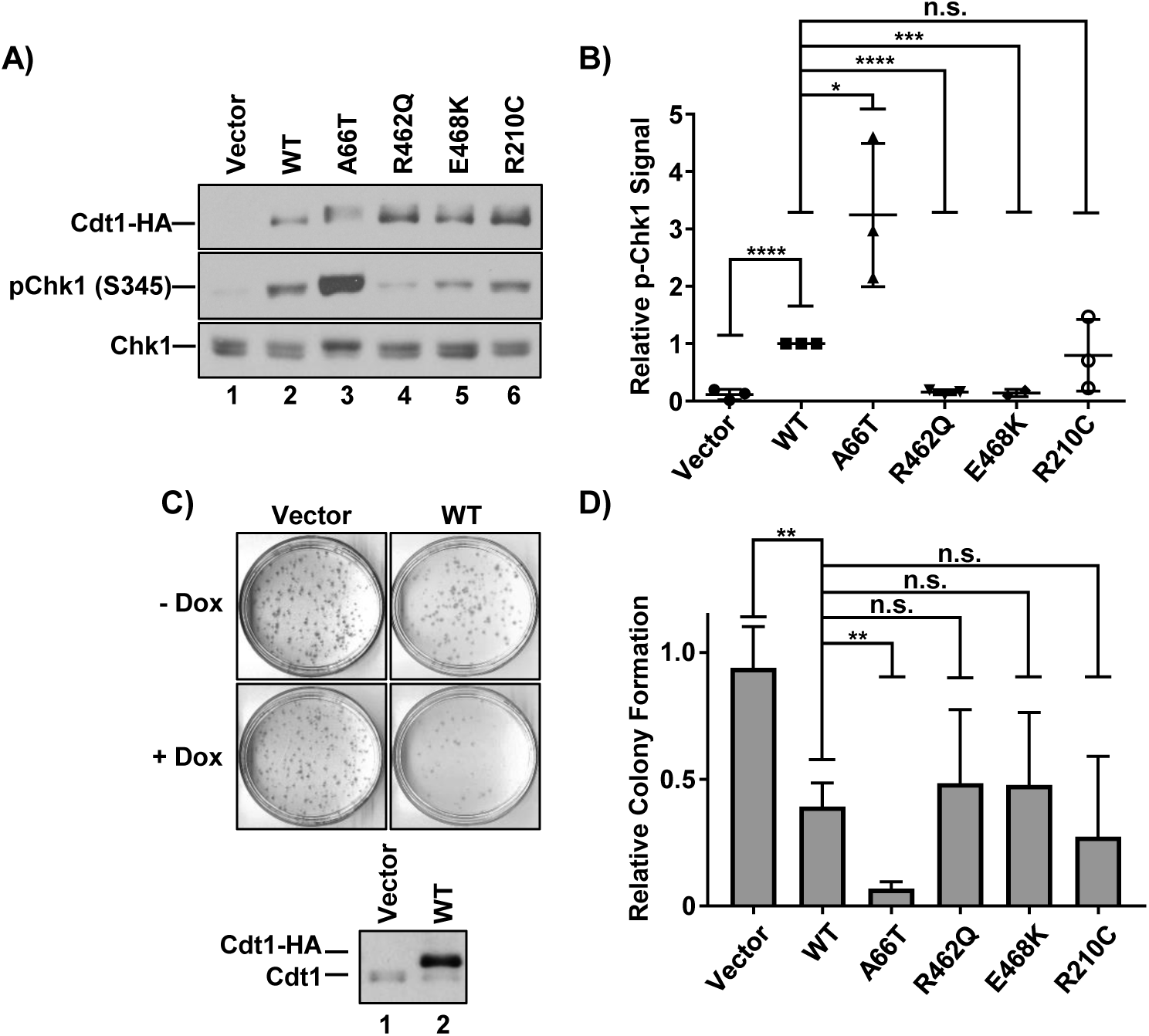
DNA damage and cell proliferation defects from Cdt1 variant overproduction. *(A)* Immunoblot of HA-tagged Cdt1 (anti-HA antibody), pChk1 (S345) and total Chk1 in U2OS cells grown in 1 µg/mL dox for 48 hours. *(B)* Graph of pChk1 (S345) induction normalized to WT Cdt1. Bars represent mean and standard deviation of three biological replicates. **** = p value <0.0001; * = p value <0.05; n.s. = not significantly different. *(C, top)* Representative vector and WT Cdt1 control colony forming assays. Cells were plated at low density in the presence or absence of 1 µg/mL doxycycline (dox) and grown for ~10 days. (*C, bottom*) A technical replicate plate was harvested after 72 hours to assay for ectopic Cdt1 expression by immunoblotting with anti-Cdt1 antibody. *(D)* Relative colony formation normalized within each experiment to the vector control; values represent at least three biological replicates. Bars represent mean and standard deviation. ** = p value <0.005; n.s. = not significantly different.

Extensive re-replication, replication stress, and DNA damage can impair cell proliferation (Li and Jin 2010; Truong and Wu 2011). As a measure of the ability of each of the Cdt1 variants to impact proliferation, we plated each cell line in either high doxycycline or no doxycycline as a control and assessed colony-formation over 10 days. Cdt1-WT overexpression strongly blocked colony formation (Fig. 2C, D). Cdt1 variants that were least active for inducing re-replication were also somewhat less capable of impairing colony formation (although these differences were not significantly different from Cdt1-WT, Fig. 2D). Strikingly however, A66T which was hyperactive for re-replication was also even more toxic than WT Cdt1 in this assay (Fig. 2D).

### Comparative functional analysis of MCM loading

Given that most Meier-Gorlin mutations affect genes encoding essential origin licensing proteins (Cdt1, Cdc6, ORC, etc.), we hypothesized that the defects associated with Cdt1 hypomorphic variants are primarily related to MCM loading. To test this idea directly, we induced expression of the Cdt1 variants in asynchronously growing cells with low doxycycline to approximately match endogenous Cdt1 levels. We simultaneously depleted endogenous Cdt1 using an siRNA; the ectopic Cdt1 expression constructs bear synonymous mutations at the siRNA binding site and are thus resistant to depletion (Fig. 3D). We then pulse labeled the cells with EdU for 30 minutes prior to harvesting and extracting cells to release soluble MCM complexes, followed by fixation to retain loaded MCM complexes. We probed the extracted cells for Mcm2 as a marker of the MCM_2-7_ complex, stained for total DNA content, detected EdU incorporation, and analyzed the samples by flow cytometry (see Materials and Methods). We previously validated this assay for quantifying MCM loading rates in asynchronously proliferating individual cells (Matson et al. 2017). Figure 3A shows flow cytometry profiles of extracted cells with chromatin-bound MCM on the y-axis and DNA content on the x-axis. MCM^Bound^-positive/ EdU-negative G1 cells are shown in blue, MCM^Bound^-positive/ EdU-positive cells are shown in orange, and MCM^Bound^-negative/ EdU-negative cells are shown in grey. Using these analytical flow cytometry profiles, we isolated the G1 phase MCM positive cells *in silico* and plotted these data in histogram form as a measure of licensing activity (Fig. 3B). In previous work, we demonstrated that these histograms reveal both the total amount of MCM loaded per cell, and the rate of MCM loading within G1 phase (Matson et al. 2017).

**Figure 3.**
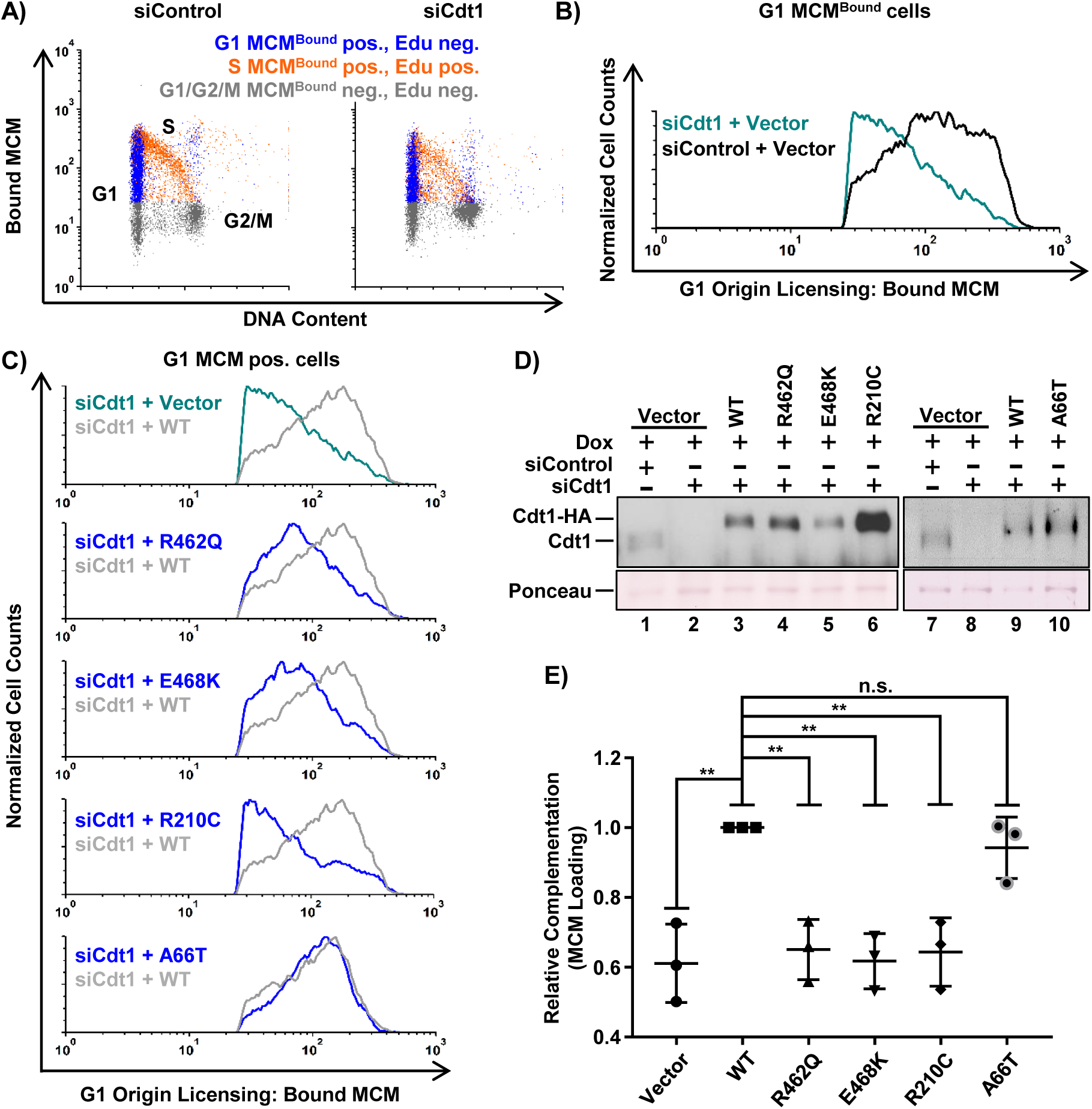
Functional analysis of MCM loading. *(A)* Analytical flow cytometry profiles of chromatin bound MCM in U2OS cells treated with 100 nM control siRNA (left) or Cdt1 siRNA (right). Cells were pulse labeled with 10 µM EdU for 30 minutes prior to harvesting and extraction of soluble MCM. Bound (unextracted) MCM was detected with anti-MCM2 antibody, and cells were stained with DAPI for total DNA content. Blue: MCM^Bound^ positive, EdU negative, G1 DNA content, Orange: EdU positive, MCM^Bound^ positive, Grey: EdU negative and MCM^Bound^ negative. *(B)* Histograms of the G1 MCM^Bound^ positive, EdU negative (i.e. blue in A) cells from both samples in *A*. Bound MCM on the x-axis and normalized cell counts on the y-axis (counts normalized to siControl). *(C)* Histograms of G1 MCM^Bound^ positive cells depleted of endogenous Cdt1 and expressing each Cdt1 variant compared to WT Cdt1 as in B. siRNA transfected cells were cultured in 0.002-0.006 µg/ml doxycycline for 72 hrs prior to EdU labeling and processing as in A. *(D)* Immunoblot of endogenous and ectopic Cdt1 from *C* detected with anti-Cdt1 antibody. *(E)* Complementation of G1 MCM loading normalized to WT Cdt1. Mean MCM^Bound^ loading intensity of each variant was divided by the mean MCM loading intensity of WT Cdt1 within each experiment. Bars represent mean and standard deviation of three biological replicates. ** = p value <0.005; n.s. = not significantly different.

As expected, Cdt1 depletion without ectopic Cdt1 expression resulted in defective MCM chromatin loading (Fig. 3B, green trace, quantified in 3E), but expression of the epitope-tagged Cdt1-WT fully complemented this MCM loading defect (Fig. 3C, compare grey and green traces). On the other hand, the R462Q and E468K variants were significantly impaired for MCM loading (Fig. 3C and 3E). Strikingly, the R210C variant was also significantly impaired for MCM loading even when it accumulated to higher levels than Cdt1-WT (Fig. 3C, D lane 6). Based on these complementation assays, we interpret the relative activity of the hypomorphs as *R462Q* =*E468K* > *R210C* (i.e. Cdt1-R210C is the weakest for G1 MCM loading when expressed at normal levels) whereas R210C is more active for inducing re-replication than R462Q and E468K when overproduced (Fig. 1).

Unlike the hypomorphic alleles, Cdt1-A66T showed no MCM loading defect in G1, which is consistent with the idea that this variant is not a loss-of-function allele (Fig. 3C and 3E). Interestingly, cells expressing Cdt1-A66T accumulated re-replicated DNA more than Cdt1-WT expressing cells even at these lower expression levels (Fig. 3D; Supplemental Fig. S3). This result suggested Cdt1-A66T hyperactivity arose from origin licensing problems outside of G1 phase, at times when Cdt1 must be tightly regulated to prevent illegitimate origin licensing.

### Comparative analysis of MCM binding

We considered that the functional effects of Cdt1 mutations may be linked to key protein-protein interactions, specifically Cdt1-MCM binding. To test this notion, we immunoprecipitated WT or variant Cdt1 using the HA epitope tag and probed for MCM_2-7_ interaction using Mcm2 as a marker of the complex. Previous studies have mapped the Cdt1-MCM binding domain to its C-terminus, and some mutations in this domain of metazoan Cdt1 impair binding to a partial MCM complex (Ferenbach et al. 2005; Jee et al. 2010). Not surprisingly then, the two hypomorphic variants located in the C-terminal domain of Cdt1, Cdt1-R462Q and Cdt1-E468K, were consistently impaired for MCM co-immunoprecipitation (Fig. 4A, B). On the other hand, Cdt1-R210 is located in the middle domain of Cdt1 and not in the previously described Cdt1-MCM binding domain. Like Cdt1-R462Q and Cdt1-E468K, Cdt1-R210C was also impaired for MCM interaction (Fig. 4C). This result suggests the existence of an MCM binding domain in Cdt1 distinct from the C-terminal domain.

**Figure 4.**
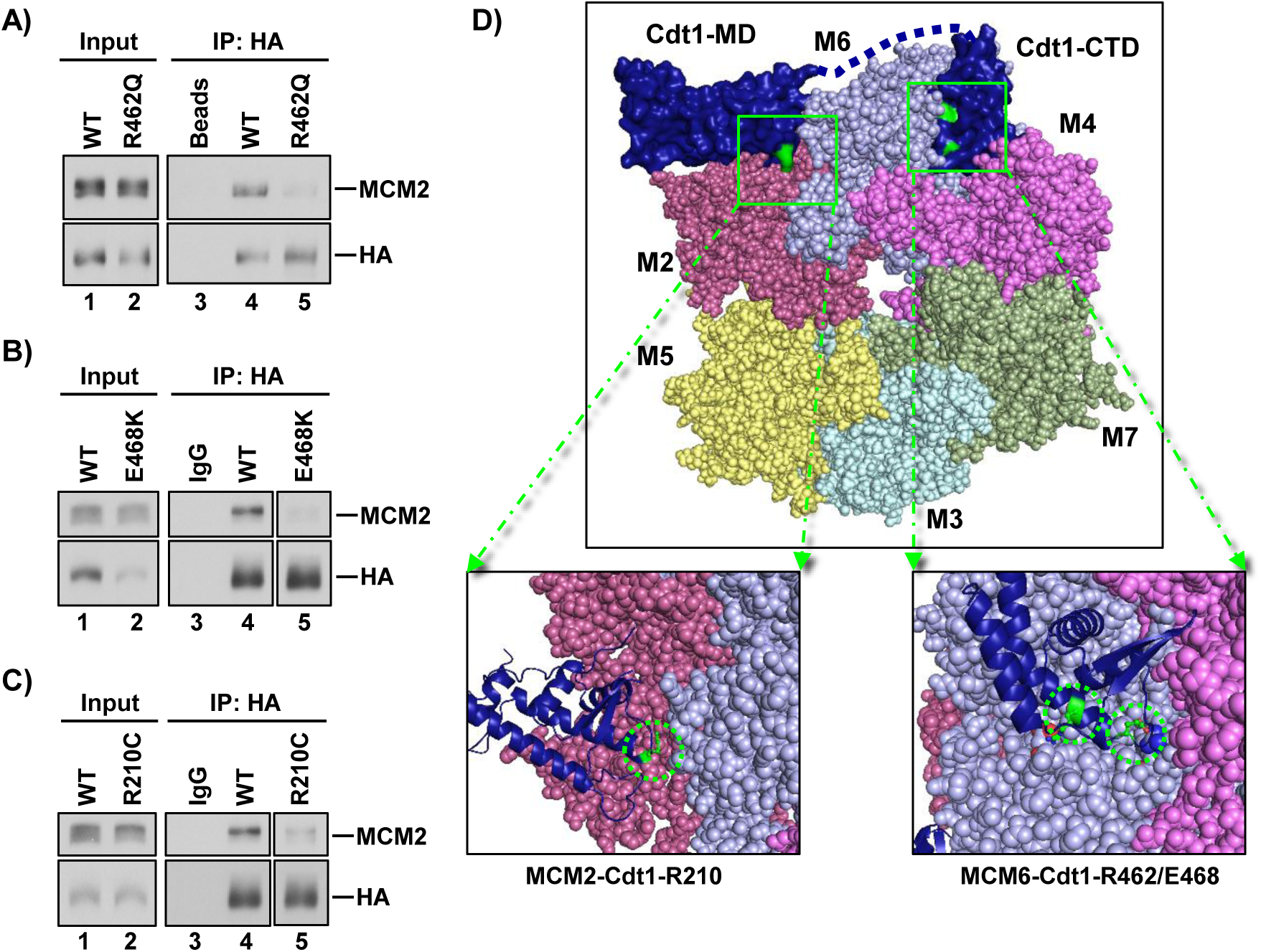
Relative MCM binding. (A-C) WT and Cdt1 variants were transiently expressed in HEK 293T cells and immunoprecipitated using the HA epitope tag. Portions (2%) of whole cell lysates and bound proteins were probed for HA-Cdt1 (anti-HA antibody) and for MCM2 as a marker of the MCM complex; spliced images are from the same immunoblot and same exposure. (D, top) Homology model of the human MCM_2-7_-Cdt1 complex. The yeast OCCM structure (PDB ID: 5UDB) was used as a template to model the human MCM_2-7_ complex;> numbers indicate individual MCM subunits, and colors are similar to Yuan *et al.*. The structure of the human C-terminal Cdt1 winged helix, “Cdt1-CTD” (PDB ID: 2WVR) and mouse Cdt1 central/middle domain “Cdt1-MD” (PDB ID: 3A4C) were used to model hCdt1-hMCM interactions. (D, bottom left) Magnified view of the proposed interacting surfaces with R210 highlighted in green. (D, bottom right) Magnified view of the proposed interacting surfaces with R462 and E468 highlighted in green.

Structures of budding yeast Cdt1 in complex with other licensing proteins have recently been reported, and although two domains of mammalian Cdt1 orthologs have been structurally characterized (Frigola et al. 2017; Yuan et al. 2017; Zhai et al. 2017), there is no complete structure of metazoan Cdt1 available. Therefore, to visualize the locations of these mutations relative to the human MCM_2-7_ complex, we generated a homology model using publicly available crystal, cryo-EM, and NMR structures of Cdt1-MCM_2-7_ complexes. We derived our model using the recent cryo-EM yeast ORC_1-6_-Cdc6-Cdt1-MCM_2-7_ (OCCM) complex (Yuan et al. 2017) as a template for modeling the human MCM complex (see Materials and Methods). We superimposed structures for the human Cdt1 C-terminal domain (De Marco et al. 2009) and mouse C-terminal (Khayrutdinov et al. 2009) on the OCCM structure to model the mammalian Cdt1-MCM_2-7_ interaction (Fig. 4D, top). The Cdt1 C-terminal domain adopts a winged helix fold of the type predicted to mediated protein-protein rather than protein-DNA interactions (Khayrutdinov et al. 2009). In this model, residues R462 and E468 are at the interface between the Cdt1 C-terminal domain and the Mcm6 subunit of the MCM_2-7_ heterohexamer (Fig. 4D, bottom right). Mutating these residues likely disrupts the binding surface between Cdt1 and Mcm6 resulting in defective Cdt1-MCM interactions.

We also consistently observed weak binding between Cdt1-R210C and MCM (Fig. 4C), but Cdt1 R210 is not in the C-terminal MCM binding domain. Our homology model positions this residue near the Mcm2 subunit of the MCM_2-7_ heterohexamer (Fig. 4D, bottom left). This proposed interface is distinct from the suggested interface between the Cdt1 C-terminal domain and Mcm6. In addition, cryo-EM structures of the yeast Cdt1-MCM_2-7_ complex predict multiple contact points between Cdt1 and the MCM_2-7_ complex (Sun et al. 2013; Yuan et al. 2017). The functional defects of Cdt1-R210C, Cdt1-R462Q, and Cdt1-E468K in cells support the notion that Cdt1 requires multiple binding interfaces with the MCM_2-7_ complex. Our functional and interaction analysis suggests that disrupting either of these interfaces is sufficient to impair both MCM_2-7_ binding and MCM_2-7_ loading.

### Cdt1-A66T impairs Cyclin A and Skp2 binding but does not stabilize Cdt1 in S phase

The unexpected gain-of-function phenotype of the A66T dwarfism-associated variant prompted us to explore this variant in more detail. Based on the close proximity of A66 to a previously defined Cyclin/CDK binding motif (“Cy motif”) at positions 68-70, we hypothesized that A66T perturbs the Cdt1-Cyclin/CDK interaction (Fig. 5A). To test this idea, we isolated Cdt1-A66T, Cdt1-WT, and Cdt1-Cy, a *bona fide* mutational disruption in the Cy motif (alanines at positions 68, 69, and 70) from cell lysates using the C-terminal tag and probed the complexes for Cyclin A; previous studies identified Cyclin A as the primary cyclin that interacts with the Cdt1 Cy motif (Sugimoto et al. 2004). Cdt1-WT bound Cyclin A by this assay, but the Cy motif mutant did not (Fig. 5B). Interestingly, Cdt1-A66T bound Cyclin A very poorly relative to Cdt1-WT and only slightly better than the Cy motif mutant (Fig. 5B, compare lane 7 to lanes 6 and 8).

**Figure 5.**
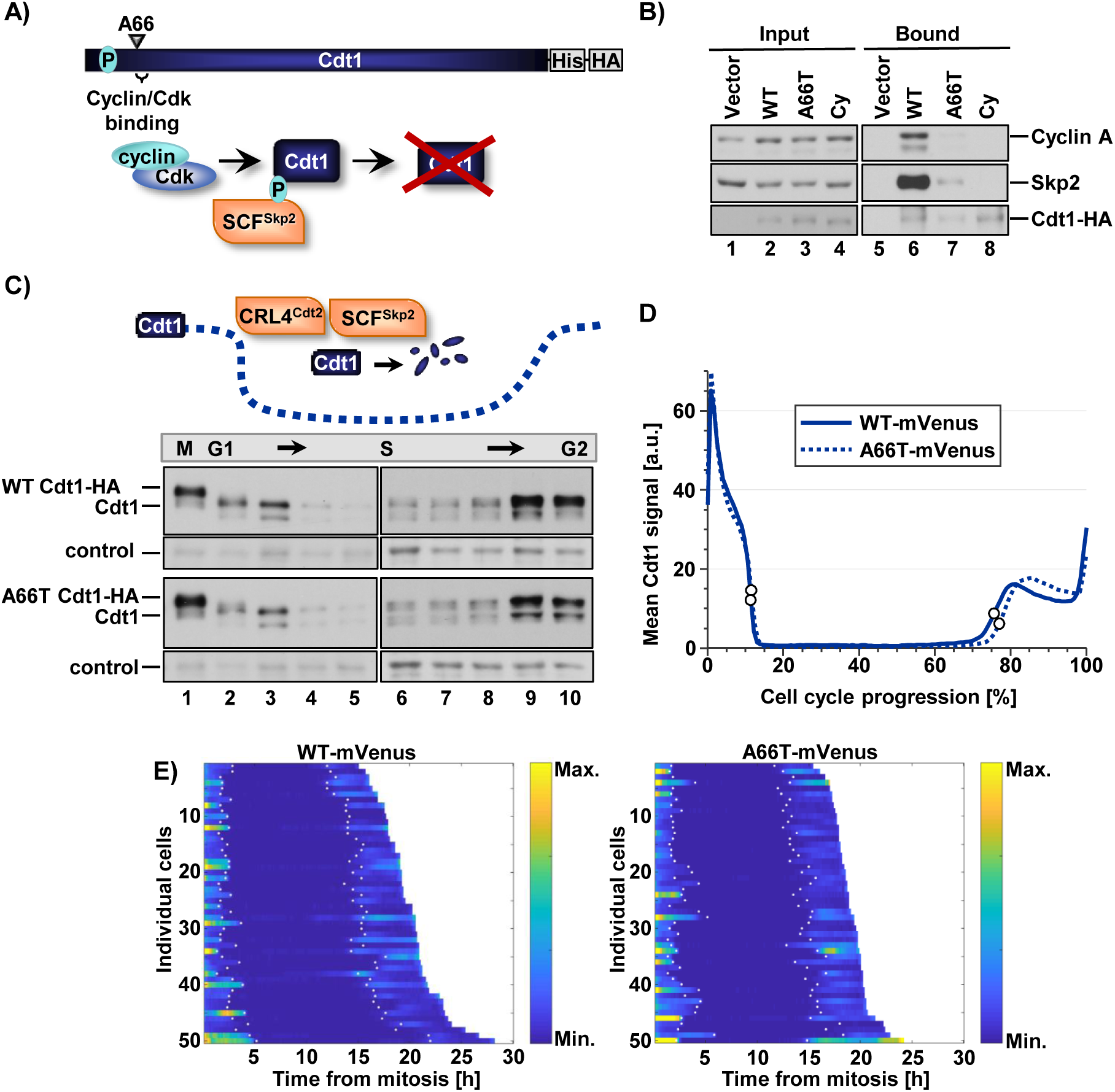
Cdt1-A66T impairs Cyclin A and Skp2 binding but does not stabilize Cdt1 in S phase. *(A)* Illustration of SCF^Skp2^-dependent degradation of WT Cdt1 via CDK-mediated phosphorylation at threonine 29. *(B)* Cells were cultured in 1 µg/mL doxycycline (high) for 18 hrs, lysed, and incubated with nickel-agarose to retrieve His-tagged Cdt1. Portions of whole cell lysates (2%) and bound complexes were probed for the indicated proteins (bottom panel anti-HA antibody). *(C, top)* Illustration of Cdt1 degradation and accumulation during the cell cycle. *(C, bottom)* U2OS cells expressing Cdt1-WT and Cdt1-A66T were synchronized by double-thymidine/nocodazole (lanes 1-6) or double-thymidine (lanes 7-12) block and released into fresh medium. Time points were taken after release and analyzed by immunoblotting with anti-Cdt1 antibody. A non-specific band serves as a loading control. *(D)* The intensity of WT or A66T Cdt1-Venus expressed in U2OS cells imaged during asynchronous proliferation every 10 minutes. Traces are the average Venus intensity in arbitrary units from mitosis (0%) to mitosis (100% cell cycle progression); n= 50 cells. White circles denote the beginning and end of S phase as determined by the localization of stably co-expressed fluorescently tagged PCNA. *(E)* Heat map of fluorescence intensity of Cdt1 WT-Venus (*left*) and Cdt1-A66T-Venus (*right*) in 50 randomly selected U2OS cells. Maps from individual cells are arranged according to the duration of the cell cycles, colors indicate differences in fluorescence levels. White dots in each track denote the beginning and end of S phase as determined by the localization of stably co-expressed fluorescently tagged PCNA.

**Figure 6.**
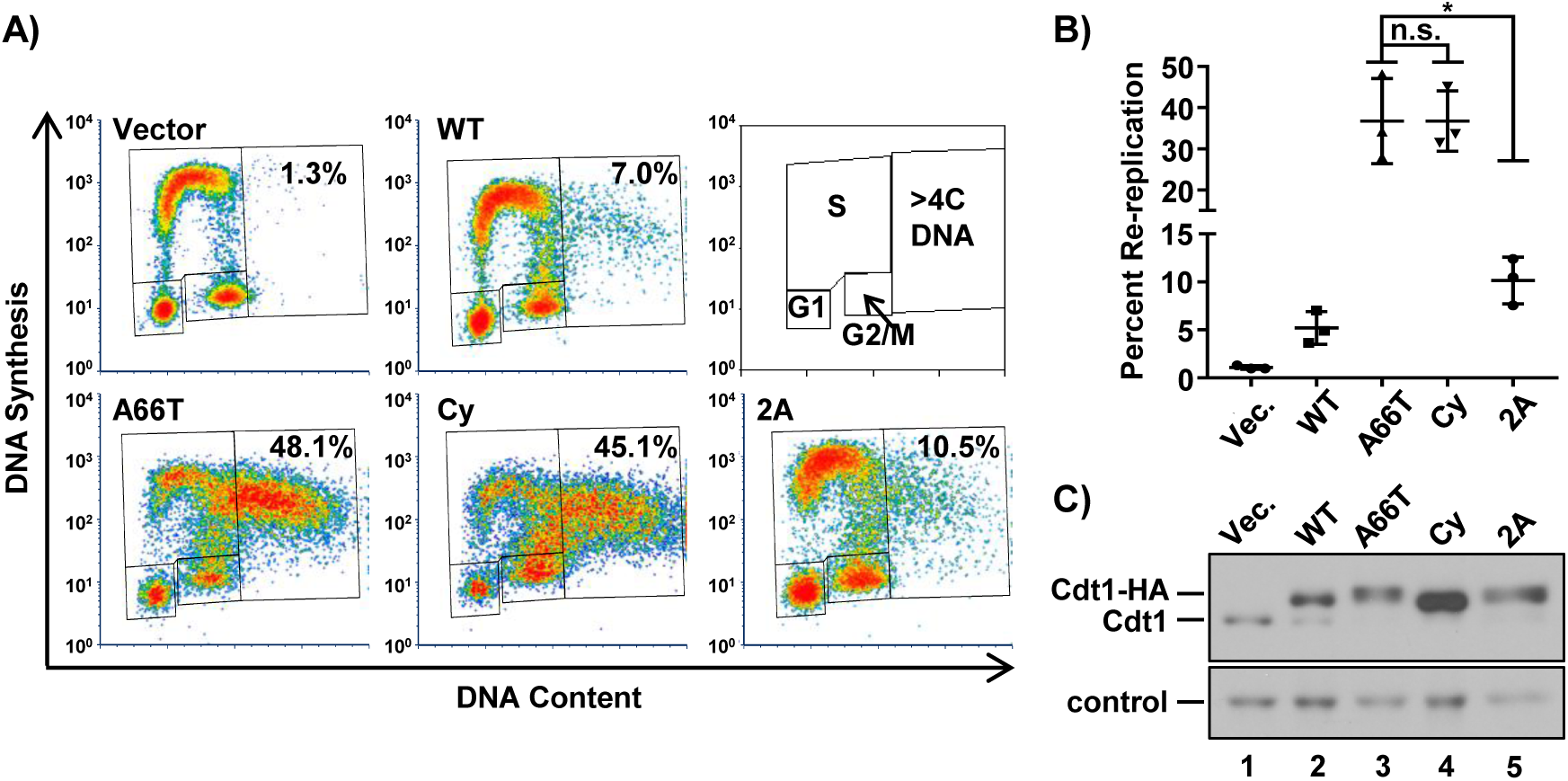
CDK-Cdt1 binding suppresses re-replication independently of the Cdt1 phosphodegron. *(A)* Analytical flow cytometry profiles of cells treated with 1 µg/mL doxycycline for 48 hrs and analyzed as in Figure 1. Cy: Cyclin/CDK binding motif mutant; 2A: Cdt1 T29A, S31A. *(B)* The percentage of cells with >4C DNA content in at least 3 biological replicates. Bars represent mean and standard deviation. * = p value <0.05; n.s. = not significantly different. *(C)* HA-tagged Cdt1 was detected by immunoblotting whole cell lysates from *A* with anti-Cdt1 antibody; a non-specific band serves as a loading control; Vec. – Vector.

The consequences of Cyclin A binding to Cdt1 have been linked to CDK-mediated Cdt1 phosphorylation at T29 (Takeda et al. 2005). Cdt1 phosphorylation at T29 creates a binding site for the Skp2 substrate adapter of the SCF E3 ubiquitin ligase, SCF^Skp2^ (Fig. 5A). We therefore tested if the weak Cyclin A binding by Cdt1-A66T also resulted in weak Skp2 binding; indeed, Cdt1-A66T bound very poorly to Skp2 compared to Cdt1-WT and slightly better than the Cdt1-Cy variant (Fig. 5B, lanes 6-8). For this reason, we tested the stability of Cdt1-A66T relative to Cdt1-WT during S phase. We synchronized cells in mitosis and released them to progress from G1 into S phase taking time points until mid-S. We also blocked cells in early S phase and released them to progress into G2, then monitored endogenous and ectopic Cdt1 by immunoblotting. We found no detectable differences between Cdt1-A66T and Cdt1-WT or endogenous Cdt1 in degradation in early S or in Cdt1 re-accumulation as S phase ends (Fig. 5C). SCF^Skp2^-mediated Cdt1 ubiquitylation cooperates with a second E3 ubiquitin ligase, CRL4^Cdt2^, to destroy Cdt1 during S phase (Abbas and Dutta 2011; Havens and Walter 2011) (Fig. 5C illustration), and this targeting does not require CDK-mediated Cdt1 phosphorylation (Arias and Walter 2005). CRL4^Cdt2^ ubiquitylates Cdt1 in both S phase and after DNA damage (Arias and Walter 2006; Jin et al. 2006; Havens and Walter 2009). We observed no effect of this mutation on Cdt1 degradation after UV irradiation (Supplemental Fig. S4), thus the A66T change has no effect on CRL4^Cdt2^ targeting.

Nonetheless, we considered that Cdt1-A66T could be slightly more stable at specific cell cycle times or in other settings in a manner that increases the likelihood of origin re-licensing and subsequent re-replication. We therefore added a C-terminal fluorescent tag to both Cdt1-WT and Cdt1-A66T (Supplemental Fig. S5) and carried out live cell imaging of asynchronously proliferating U2OS cells after doxycycline-induced expression. We tracked individual cells with similar maximum fluorescence intensities for both Cdt1-WT and Cdt1-A66T and plotted both the mean (Fig. 5D) and the intensity values of 50 individual proliferating cells (Fig. 5E). Importantly, we observed no statistically significant differences in the dynamics of Cdt1 degradation and re-accumulation during the cell cycle for those cells that successfully divided. Moreover those A66T-expressing cells that arrested with large nuclei (presumably from re-replication (Melixetian et al. 2004)) had normal degradation and accumulation in the S phase prior to the arrest (data not shown).

Cdt1-A66T is largely impaired for SCF^Skp2^ binding, but there were no detectable consequences for Cdt1 stability since Cdt1-A66T levels are still subject to CRL4^Cdt2^ control. Nonetheless, Cdt1-A66T is a potent re-replication inducer. We thus considered that the mutation has consequences for Cdt1 activity beyond phosphorylation at T29. To test that idea directly, we expressed a Cdt1 phosphorylation site mutant in which T29 is converted to unphosphorylatable alanine. Cdt1 is also phosphorylated at S31 (Hornbeck et al. 2015), and although this phosphorylation has minimal impact on Skp2 binding compared to T29 phosphorylation (Takeda et al. 2005), we also converted S31 to alanine to avoid possible compensatory effects at this position; this double alanine mutant is “Cdt1-2A.” If the primary effect of Cdt1-A66T is to prevent phosphorylation at T29 (and S31), then we predicted that the phenotypes of cells overexpressing Cdt1-A66T, the Cdt1-Cy motif mutant, and Cdt1-2A should be similar, since each alteration blocks CDK-mediated T29 phosphorylation. We compared the DNA re-replication activity induced by overproducing each of these Cdt1 variants. Strikingly both Cdt1-A66T and Cdt1-Cy induced significantly more re-replication than Cdt1-2A did (Fig. 6A and 6B). In these longer-expression experiments, Cdt1-Cy accumulates to higher levels than WT (Fig. 6C), though we note that in shorter experiments such as Fig. 5B, Cdt1-Cy levels are similar to Cdt1-WT and Cdt1-A66T. This hyper-accumulation may be a consequence of cell cycle phase distribution from long-term expression (Supplemental Fig. S6) and/or of the apparent complete defect in SCF^Skp2^ binding. Nonetheless, Cdt1-A66T and Cdt1-2A routinely accumulate to similar levels (Fig. 6C, compare lanes 3 and 5), yet Cdt1-A66T induces significantly more re-replication than Cdt1-2A does (Fig. 6A and 6B). We thus conclude that Cdt1-A66T disrupts Cyclin A binding as a near-mimic of the engineered Cdt1-Cy motif mutant, and that Cyclin A binding to Cdt1 negatively regulates Cdt1 function by at least one mechanism that is independent of simply creating a phosphodegron binding site for the SCF^Skp2^ E3 ubiquitin ligase.

## DISCUSSION

In this study, we analyzed naturally arising mutations in Cdt1 and demonstrate that hypo-and hypermorphic variants cause defects in cell proliferation through distinct molecular mechanisms. Specifically, Cdt1 mutations found in humans afflicted with Meier-Gorlin syndrome result in either Cdt1-MCM binding or Cyclin/CDK binding defects. Both scenarios lead to proliferation defects arising from problems in DNA replication that can be attributed to perturbed Cdt1 activity (Fig. 7).

**Figure 7.**
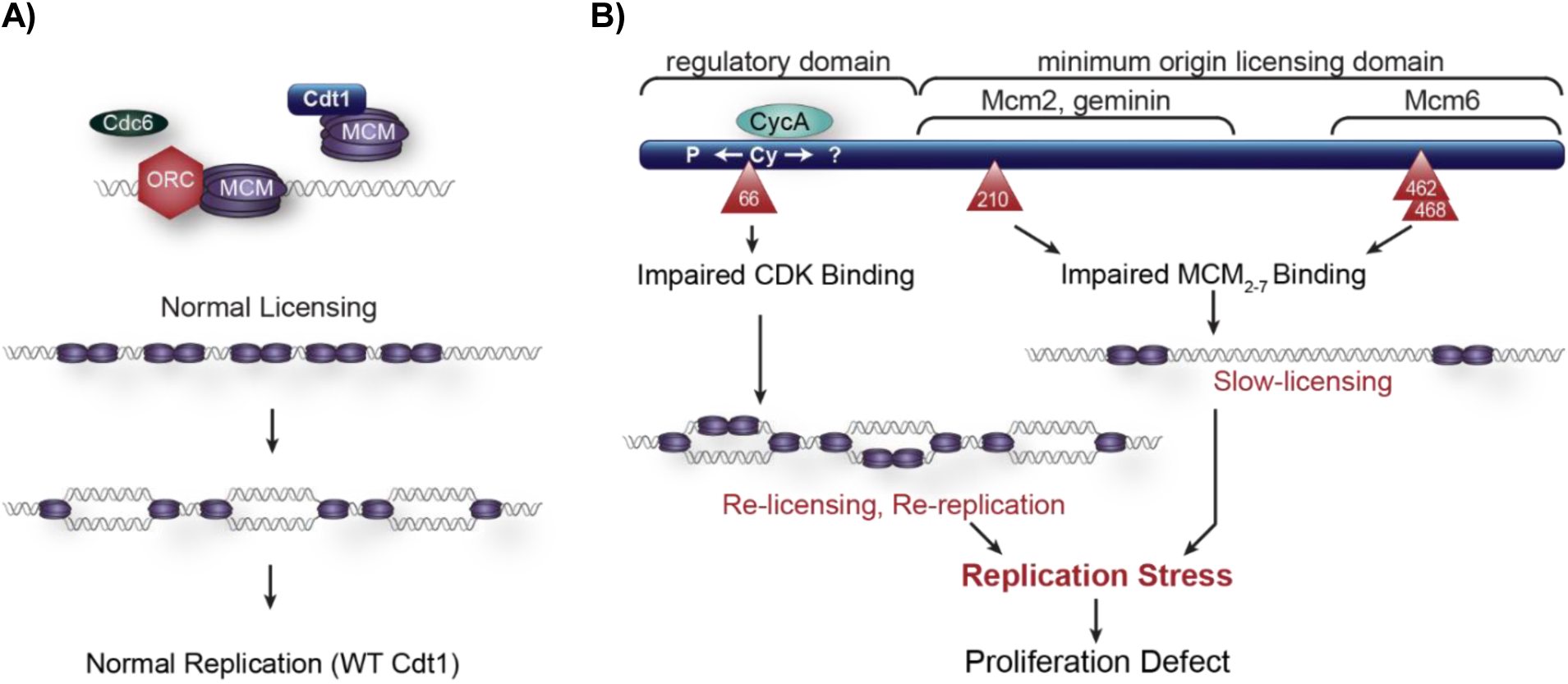
Model of cell proliferation defects from both hypomorphic or hypermporphic Cdt1 variants. *(A)* WT Cdt1 supports normal MCM loading/origin licensing and normal DNA replication in S phase. *(B)* The A66T variant is impaired for CDK-mediated repression resulting in re-licensing and re-replication. The R210C, R462Q, and E468K variants are impaired for MCM2-7 binding in G1 phase leading to insufficient origin licensing. Both scenarios ultimately lead to DNA replication stress and proliferation defects.

### Hypomorphic Alleles

Meier-Gorlin (MG) syndrome is a form of primordial dwarfism characterized by growth retardation beginning *in utero* and continuing throughout adolescence. Based on the patients’ phenotypes, we hypothesized that all MG *Cdt1* alleles are hypomorphic. Indeed, two of the alleles analyzed here, including *Cdt1-R462Q* which was present in 9 of the 11 MG patients with Cdt1 mutations reported thus far, are hypomorphic for Cdt1 function (Bicknell et al. 2011a; Bicknell et al. 2011b; Guernsey et al. 2011; de Munnik et al. 2012). Q117 is poorly conserved among Cdt1 sequences suggesting that it is not critical for Cdt1 function. Based on its apparently normal ability to induce re-replication and the poor conservation of Q117, we infer that Cdt1-Q117H is hypomorphic for Cdt1 <u>expression</u> in the MG patient rather than <u>function</u> – possibly from inefficient mRNA splicing (Bicknell et al. 2011a). The mutation may reduce overall Cdt1 expression *in vivo* rather than impact Cdt1 activity *per se.* R453 is buried in the winged-helix domain core of the human Cdt1 C-terminal domain (Khayrutdinov et al. 2009; Jee et al. 2010). Introducing a bulky aromatic tryptophan may globally disrupt folding rather than alter Cdt1 interactions or function.

The two functionally hypomorphic MG alleles in this study, *Cdt1-R462Q* and *Cdt1-E468K*, encode substitutions of conserved solvent-exposed amino acids in the C-terminal Cdt1 winged helix domain (Khayrutdinov et al. 2009; Jee et al. 2010). By analytical flow cytometry, we found that these variants support slower MCM loading relative to WT Cdt1. The hypomorphic nature of these alleles could induce slow proliferation by lengthening G1 phase through an origin licensing checkpoint and/or cells may arrest from under-replication during S phase itself. Our homology model places R462 and E468 at the interface between the Cdt1 C-terminal domain and the Mcm6 subunit of the MCM_2-7_ heterohexamer, close to the Mcm6-Mcm4 interface. Thus, mutations in this region understandably impair MCM binding.

*Cdt1-R210C* is orthologous to a mutation in the *Drosophila melanogaster Cdt1* gene, Double-Parked (Dup). Whittaker et al. characterized this variant as hypomorphic resulting in DNA replication defects and female sterility (Whittaker et al. 2000). Previous studies reported that this Cdt1 variant supported less DNA synthesis *in vitro* (De Marco et al. 2009), and had a modest effect on migration of a Cdt1-MCM_2-7_ complex by native gel electrophoresis (You et al. 2016). We found that this variant has impaired Cdt1-MCM_2-7_ binding by co-immunoprecipitation from human cell lysates. This variant supports only slow origin licensing that is nearly as slow as the two dwarfism hypomorphic alleles in the C-terminal domain. The similarity in both cellular and molecular phenotypes of *Cdt1-R462Q*, *Cdt1-E468K*, and *Cdt1-R210C* suggests that both the central domain and C-terminal domain are equally important for Cdt1-MCM binding.

Jee et al. suggested the existence of cooperation between the central domain of Cdt1 and the C-terminal domain in origin licensing (Jee et al. 2010). Yanagi et al. found that a fragment of murine Cdt1 including the central domain but lacking the Cdt1 C-terminal domain can associate with a subcomplex of three subunits, MCM4/6/7 (Yanagi et al. 2002). The notion of multiple contacts between Cdt1 and MCM_2-7_ is consistent with recent structural and functional analysis of yeast Cdt1-MCM_2-7_ in which Cdt1 serves as a brace to keep the MCM_2-7_ ring open during MCM loading (Frigola et al. 2017). In this model, Cdt1 must engage the MCM_2-7_ complex at two distinct points to maintain the Mcm2/Mcm5 “gate” open for DNA entry during MCM loading.

The existence of a second MCM_2-7_ binding site in the central region of Cdt1 sheds light on the mechanism of Cdt1 inhibition by the origin licensing inhibitor protein, Geminin. Previous studies have shown that Geminin inhibits Cdt1 by blocking its interaction with the MCM complex (Yanagi et al. 2002; Cook et al. 2004). The molecular mechanism of that interference cannot be easily explained if the only place MCM binds Cdt1 is the C-terminal domain. The co-crystal structure of Cdt1 in complex with Geminin includes only the central region of Cdt1 (including R210) and not the C-terminal domain (Lee et al. 2004). If the central domain is also essential for MCM_2-7_ binding, then we postulate that it is only this interaction that geminin targets. Moreover, interfering with binding either interface is sufficient to impair overall MCM_2-7_ binding and therefore, MCM_2-7_ loading.

### Dwarfism Hypermorphic Allele

Cdt1 is tightly regulated throughout the cell cycle to ensure once-and-only once DNA replication. One of the mechanisms to restrict Cdt1 activity outside of G1 phase and avoid re-replication is ubiquitin-mediated proteolysis. This process is carried out by two E3 ubiquitin ligases, CRL4^Cdt2^ and SCF^Skp2^ (Nishitani et al. 2006) (Fig. 5C, top). CRL4^Cdt2^ relies on chromatin-bound PCNA to ubiquitylate its PIP-degron containing substrates (Arias and Walter 2006; Jin et al. 2006; Havens and Walter 2011). On the other hand, ubiquitylation of Cdt1 by SCF^Skp2^ is dependent on Cyclin/CDK phosphorylation generating a phosphodegron which is recognized by the Skp2 adapter subunit (Li et al. 2003; Liu et al. 2004). Given the proximity of the A66T mutation to the Cyclin/CDK binding motif coupled with the defect in Cyclin A and Skp2 binding, we first reasoned the hyperactivity of this variant was due to increased protein stability. Cdt1-A66T is not more stable than WT Cdt1 however, so impaired degradation does not explain this variant’s phenotype.

Our comparison of Cdt1-A66T, an engineered null for Cyclin/CDK binding (Cdt1-Cy), and a variant that can bind Cyclin but cannot generate a phosphodegron (Cdt1-2A) directly demonstrated that Cyclin A-dependent regulation of Cdt1 involves more than just degradation, because mutating the phosphodegron had less impact than mutating the Cyclin/CDK binding site (Fig. 6). We thus postulate that Cyclin/CDK also inhibits Cdt1 by non-degradation mechanisms. Coulombe *et al*. described a negative-regulatory PEST domain (a.a. 74-108) in mammalian Cdt1 that contains multiple candidate CDK phosphorylation sites (Coulombe et al. 2013). This domain functions independently of either Geminin or the E3 ubiquitin ligase system. Deleting the PEST domain induced DNA re-replication similar to Cdt1-A66T. Cyclin/CDK may phosphorylate any of the other CDK target residues – either in the PEST domain or elsewhere - which could inhibit Cdt1 activity. A total of 20 candidate CDK phosphorylation sites have been detected in human Cdt1 by mass spectrometry, and only 7 of these have been functionally tested so far (Pozo and Cook 2016). Given the apparent efficient interaction of Cdt1 with Cyclin A/CDK, it is also possible that Cyclin binding itself inhibits Cdt1 activity independently of phosphoregulation.

It is surprising that mutational alterations that lead to similar phenotypes in Meier-Gorlin Syndrome dwarfism patients behave differently at the molecular level with respect to Cdt1. In the case of Cdt1-R462Q and Cdt1-E468K, impaired Cdt1-MCM interactions can lead to slower G1 progression due to slow origin licensing. On the other hand, Cdt1-A66T dysregulation by Cyclin/CDK results in replication stress, which can also lead to proliferation failure (Fig. 7). The ultimate outcome in either case however is impaired overall growth. Continual improvements in our understanding of the molecular mechanisms governing origin licensing are essential to link processes of cell proliferation, genome stability, and development.

## MATERIALS AND METHODS

### Cell culture and manipulations

U2OS Flp-in Trex (Malecki et al. 2006) cells bearing a single FRT site (gift of J. Aster) and HEK 293T cells were cultured in Dulbecco’s modified Eagle’s medium (DMEM) supplemented with 10% fetal bovine serum (FBS) and 1x penicillin/streptomycin (complete medium). To generate stable isogenic cell lines, U2OS cells were co-transfected with flippase recombinase (Flp) and a Cdt1 expression vector derived from pcDNA5/FRT/TO-Venus-Flag-Gateway (1124), a gift from Jonathon Pines (Addgene plasmid # 40999), using X-tremeGENE HP DNA transfection reagent (Roche). The Cdt1 cDNAs encode normal Cdt1 or harbor a single point mutation and a drug resistance cassette. 48 hrs post-transfection, cells were selected for resistance to either 150 µg/mL hygromycin B (Roche) or 1 µg/mL puromycin (Sigma) depending on the Cdt1 vector used. For inducible expression of Cdt1 variants, U2OS cells were treated with varying concentrations of doxycycline ranging from 0.003 µg/mL-1 µg/mL (CalBiochem) by either media exchange or adding directly into cell culture plates. For colony forming assays, U2OS cells harboring Cdt1 mutant alleles were plated at a density of ~500 cells/ 6 cm dish in the presence or absence of doxycycline. Cells were grown for 10 days, changing media every three days, and stained using 0.4% crystal violet (Fisher Scientific). Colony numbers and size were quantified using ImageJ (NIH). A technical replicate plate was harvested after 72 hours to assay for immunoblot analysis.

For G1 to S phase synchronization, U2OS cells were treated with 2.5 mM thymidine for 24 hours followed by release into complete medium containing 100 ng/mL nocodazole plus 0.05 µg/mL doxycycline for 16 hours. Cells were then harvested by mitotic shake-off and re-plated in complete medium plus 0.05 µg/mL doxycycline for each time point. For S to G2/M phase synchronization, U2OS cells were treated with 2.5 mM thymidine for 18 hours followed by release into complete medium for 8 hours. Cells were then treated with 2.5 mM thymidine plus doxycycline for 18 hours followed by release into complete medium plus doxycycline for each time point. To transiently express Cdt1 variants, HEK 293T cells were transfected with Cdt1 expression vectors using PEI Max (Sigma) according to the manufacturer’s instructions. HEK 293T cells were harvested after 16 hours post-transfection and processed for subsequent co-immunoprecipitation assays.

All cell lines were validated by STR profiling and tested mycoplasma-negative.

### Plasmids

Cdt1 mutations (Cdt1-A66T, -Q117H, -R210C, -R453W, -R462Q, -E468K) were generated by PCR-based mutagenesis from a WT Cdt1 coding sequence template. The resulting PCR products were cloned into pENTR vectors harboring the full-length Cdt1 sequence with C-terminal polyhistidine (His) and hemagglutinin (HA) epitope tags. The Cdt1-Y520X truncation was generated using Gibson Assembly (NEB) from a pENTR plasmid harboring a WT version of Cdt1 with C-terminal polyhistidine and HA epitope tags, following the manufacturer’s protocols. The pENTR-EGFP (vector control) plasmid was generated by subcloning EGFP from an EGFP bearing plasmid into pENTR via Gateway Cloning (Invitrogen). EGFP, Cdt1-WT-His-HA, Cdt1-Y520X-His-HA, Cdt1-Mutant-His-HA, Cdt1-Cy-His-HA, and Cdt1-2A-His-HA versions were transferred from pENTR plasmids into derivatives of pcDNA5/FRT/TO-Venus-Flag-Gateway (1124), harboring either hygromycin B (Roche) or puromycin (Sigma) selection cassettes, via Gateway Cloning. The mVenus tagged constructs were constructed by subcloning mVenus into the Cdt1-WT-His-HA or the Cdt1-A66T-His-HA pENTR plasmids before Gateway cloning into pcDNA5/FRT/TO-Venus-Flag-Gateway (1124).

### Flow cytometry analysis for DNA re-replication

U2OS cell lines harboring stably integrated individual Cdt1 alleles were cultured in complete medium plus doxycycline for either 48 or 72 hours. Cell were pulse labeled with 10 µM EdU (Sigma) for 30 minutes prior to harvesting by trypsinization. Approximately 20% of this suspension was reserved for subsequent immunoblotting analysis. The remaining 80% was fixed in 1 x PBS plus 4% paraformaldehyde (Sigma) at room temperature for 15 minutes. Cells were permeabilized in 1% BSA plus 0.5% triton X-100 for 15 minutes then processed for EdU detection by conjugation to Alexa Fluor 647 azide (Life Technologies) in 1 mM CuSO_4_ and 100 mM ascorbic acid; total DNA was detected by staining with 1 µg/mL DAPI (Life Technologies) in 100 µg/mL RNAse A (Sigma). Samples were analyzed on a Beckman Coulter CyAn ADP cytometer and data analyzed using FCS Express 6 (De Novo Software) software.

### MCM loading analysis by flow cytometry

U2OS cells lines harboring stably integrated individual Cdt1 alleles were plated into dishes containing a mixture of siRNA (100 nM final concentration), Dharmafect 1 (Dharmacon), and antibiotic free media plus doxycycline for 72 hours. Cells were pulse labeled with 10 µM EdU (Sigma) for 30 minutes prior to harvesting by trypsinization. Approximately 20% of this suspension was reserved for subsequent immunoblotting analysis while the remaining 80% was analyzed for bound MCM as described and validated in Matson et al. (Matson et al. 2017) and (Haland et al. 2015; Moreno et al. 2016). Briefly, cells were extracted in cold CSK buffer (10 mM Pipes pH 7.0, 300 mM sucrose, 100 mM NaCl, 3 mM MgCl_2_) supplemented with 0.5% triton X-100, protease inhibitors (0.1 mM AEBSF, 1 µg/mL pepstatin A, 1 µg/mL leupeptin, 1 µg/mL aprotinin), and phosphatase inhibitors (10 µg/mL phosvitin, 1 mM β-glycerol phosphate, 1 mM Na-orthovanadate). Cells were washed with PBS plus 1% BSA and then fixed in 4% paraformaldehyde (Sigma) followed by processing for EdU conjugation to Alexa Fluor 647 azide (Life Technologies). Bound MCM was detected by incubation with anti-MCM2 primary antibody at 1:200 dilution and anti-mouse-488 at 1:1,000 dilution at 37 °C for 1 hour. Total DNA was detected by incubation in 1 µg/mL DAPI (Life Technologies) and 100 µg/mL RNAse A (Sigma). Samples were processed on a Beckman Coulter CyAn ADP cytometer and data analyzed using FCS Express 6 (De Novo Software) software. Control samples were prepared omitting primary antibody or EdU detection to define thresholds of detection.

### Antibodies

The following antibodies were purchased from Cell Signaling Technologies: anti-pChk1 S345 (Cat# 2341), anti-Chk1 (Cat# 2345), anti-Cdt1 (Cat# 8064), anti-Skp2 (Cat# 4313). Anti-HA used for immunoblotting was purchased from Roche (Cat# 11867423001). Anti-HA used for co-immunoprecipitation was purchased from Santa Cruz Biotechnology (Cat# SC-805). Anti-Cyclin A was purchased from Santa Cruz Biotechnology (Cat# SC-596). Anti-MCM2 was purchased from BD Biosciences (San Jose, CA, Cat#610700). Anti-mouse Alexa 488 (Jackson ImmunoResearch) and Alexa 647-azide (Life Technologies) were used in flow cytometry analyses. Secondary antibodies for immunoblotting were purchased from Jackson ImmunoResearch.

### Protein-protein interaction assays

For HEK 293T co-immunoprecipitation assays, cells were transiently transfected using expression vectors harboring individual Cdt1 alleles. Cells were harvested by trypsinization, pelleted, and resuspended in co-IP Buffer (50 mM HEPES pH 7.2, 33 mM KAc, 1 mM MgCl_2_, 0.5% triton X-100, 10% glycerol) containing protease inhibitors (0.1 mM AEBSF, 10 µg/mL pepstatin A, 10 µg/mL leupeptin, 10 µg/mL aprotinin), phosphatase inhibitors (5 µg/mL phosvitin, 1 mM β-glycerol phosphate, 1 mM Na-orthovanadate), 1 mM ATP, and supplemented with 5 mM CaCl_2_ and 15 units of S7 micrococcal nuclease (Roche). Lysates were sonicated for 10 seconds on low power followed by incubation on ice for 20 minutes and clarification by centrifugation at 13,000 x g at 4°C. Supernatants were pre-cleared with Protein A-Agarose (Roche) then incubated with 1 µg antibody at 4°C overnight with rotation. Antibody-antigen complexes were collected on Protein A beads at 4°C for 1 hour with rotation. Complexes were washed 3 times rapidly with 1 mL ice-cold co-IP buffer then eluted by boiling in SDS sample buffer supplemented with 10% β-ME and 100 mM DTT for subsequent immunoblot analysis.

For polyhistidine pulldown assays, U2OS cells harboring each individual allele were plated in complete medium plus 1 µg/mL doxycycline for 16 hours, then lysed in 50 mM HEPES pH 8.0, 33 mM KAc, 117 mM NaCl, 20 mM Imidazole, 0.5% triton X-100, 10% glycerol) containing protease inhibitors (0.1 mM AEBSF, 10 µg/mL pepstatin A, 10 µg/mL leupeptin, 10 µg/mL aprotinin), phosphatase inhibitors (5 µg/mL phosvitin, 1 mM β-glycerol phosphate, 1 mM Na-orthovanadate), 1 mM ATP, 1 mM MgCl_2_, and supplemented with 5 mM CaCl_2_ and 15 units of S7 micrococcal nuclease (Roche). Clarified lysates were incubated with nickel NTA agarose (Qiagen) for 2 hours at 4°C with rotation. Beads were washed 4 times rapidly with 1 mL ice cold lysis buffer then boiled in sample buffer prior to immunoblot analysis.

### Live-cell imaging and analysis

U2OS cells stably expressing a PCNA-mTurquoise2 fusion (introduced by retroviral transduction) were plated on glass-bottom plates (Cellvis) #1.5 in FluoroBrite DMEM (Invitrogen) supplemented with FBS, L-glutamine, and penicillin/streptomycin and kept in a humidified chamber (Okolabs) at 37°C with 5% CO_2_. A Nikon Ti Eclipse inverted microscope with Plan Apochromat dry objective lenses 20x (NA 0.75), Nikon Perfect Focus System and Andor Zyla 4.2 sCMOS detector with 12 bit resolution was used for imaging. For fluorescence imaging Chroma filters were optimized for YFP spectral range - excitation: 500/20 nm, beam splitter: 515 nm and emission: 535/30 nm. Images were collected every 10 minutes using NIS-Elements AR software. No photobleaching or phototoxicity was observed in imaged cells. Expression of Cdt1 (WT/A66T) – mVenus was induced with 50 ng/ml Dox (A66T) or 100 ng/ml Dox (WT).

Image and data analysis were performed using Fiji, ImageJ NIH (Schindelin et al. 2012) software (version 1.51n) and Matlab (R2017b MathWorks). Briefly, asynchronous colonies of cells were followed in time-lapse experiments and individual cells were tracked, segmented and synchronized *in silico*. Before the analysis, images were background corrected using rolling ball subtraction. Individual cells were tracked in a user-assisted way and nuclear regions were segmented based on PCNA images. These regions of interest were used to measure Cdt1 (WT/A66T) – mVenus intensity. Cells in S phase were detected based on S-phase punctate pattern of PCNA by calculating the variance of fluorescence intensity of PCNA in a spatial scale corresponding to foci size. Cells lacking sufficient PCNA contrast to confidently detect S phase boundaries were manually removed from the analysis set. To visualize dynamics of mean Cdt1 (WT/A66T) signal throughout the cell cycle in a population of cells, the signals collected from individual cells were normalized to cell cycle type in this manner: cell cycle phases were defined for individual cells based on PCNA localization, and traces of Cdt1 intensity were linearly interpolated over the expected number of time points in each cell cycle phase (based on measurements of median cell cycle phase lengths in the population). This *in silico* alignment emphasized the sharp changes in protein abundance at the boundaries of cell cycle phases rather than the smoothing from averaging cells with different lengths of individual phases.

### Structural model of hMCM complex with hCdt1 middle domain and C-terminal winged-helix domain

The atomic resolution structure of the yeast MCM_2-7_, Cdc6, ORC_1-6_, and Cdt1 complex was determined by electron microscopy at a resolution of 3.9 Å (PDB ID 5udb (Yuan et al. 2017)). This structure was the template used for modeling the atomic structures of the human MCM (hMCM) complex as well as the interaction of human Cdt1 (hCdt1) with hMCM. MCM subunits are highly conserved during evolution, with yeast and human subunits sharing 46-50% sequence identity. Human and yeast MCM2, MCM4 and MCM6 subunits, in particular, share 50%, 47% and 47% sequence identity, respectively. Modeller v9.16 (Marti-Renom et al. 2000) was used to generate the structural models of human MCM subunits using the yeast MCM subunits (PDB ID 5udb) as a template. No modeling of hCdt1 was needed as X-ray crystallography had been used to determine the structure of the N-terminal winged helix domain of hCdt1 at a resolution of 3.3 Å (PDB ID 2wvr (De Marco et al. 2009)), while the C-terminal winged helix domain of mouse Cdt1 was determined at 1.89 Å resolution (PDB ID 3a4c (Khayrutdinov et al. 2009)). Due to the low sequence conversation between yeast and human Cdt1, the two mammalian winged helix domains were superimposed on the corresponding yeast Cdt1 winged helix domains in 5udb using the sequence-independent and structure-based dynamic programming alignment method accessed through the ‘align’ command in the PyMOL molecular vision system (The PyMOL Molecular Graphics System, Version 2.0 Schrödinger, LLC.).

### Quantification and statistical analyses

Data were analyzed using GraphPad Prism 7.0 and MATLAB and Statistics Toolbox Release 2017b, The MathWorks, Inc., Natick, Massachusetts, United States.

## ACKNOWLEDGMENTS

We thank J. Aster for the generous gift of U2OS TRT TrEX cells, and all members of the Cook lab for their insightful discussions in preparing this manuscript. This project was supported by the National Institutes of Health F31GM121073 to P.N.P., the National Science Foundation Graduate Student Fellowship DGE-1144081 to J.P.M., and by the National Institutes of Health R01GM102413, R25GM055336, and a grant from the W.M. Keck Foundation to J.G.C; T32CA009156 supported G.D.G.. The UNC Flow Cytometry Core Facility is supported in part by P30 CA016086 Cancer Core Support Grant to the UNC Lineberger Comprehensive Cancer Center.

## Author contributions

P.N.P. and J.G.C. conceived the study; P.N.P. performed and designed most of the experiments; K.M. K. and G.D.G. performed the live cell imaging and analysis; B.T. produced the structural homology data; J.P.M. performed the MCM loading analysis by flow cytometry; Y.C. performed the colony forming assay and pChk1 immunoblotting; J.P.M., Y.C., G.D.G., provided input and discussion for the manuscript; P.N.P. produced the figures, interpreted the results, analyzed the data, and wrote the manuscript with J.G.C.

## REFERENCES

Abbas T, Dutta A. 2011. CRL4Cdt2: master coordinator of cell cycle progression and genome stability. Cell Cycle 10: 241–249.

Arentson E, Faloon P, Seo J, Moon E, Studts JM, Fremont DH, Choi K. 2002. Oncogenic potential of the DNA replication licensing protein CDT1. Oncogene 21: 1150–1158.

Arias EE, Walter JC. 2005. Replication-dependent destruction of Cdt1 limits DNA replication to a single round per cell cycle in Xenopus egg extracts. Genes Dev 19: 114–126.

Arias EE, Walter JC. 2006. PCNA functions as a molecular platform to trigger Cdt1 destruction and prevent re-replication. Nat Cell Biol 8: 84–90.

Bicknell LS, Bongers EM, Leitch A, Brown S, Schoots J, Harley ME, Aftimos S, Al-Aama JY, Bober M, Brown PA et al. 2011a. Mutations in the pre-replication complex cause Meier-Gorlin syndrome. Nat Genet 43: 356–359.

Bicknell LS, Walker S, Klingseisen A, Stiff T, Leitch A, Kerzendorfer C, Martin CA, Yeyati P, Al Sanna N, Bober M et al. 2011b. Mutations in ORC1, encoding the largest subunit of the origin recognition complex, cause microcephalic primordial dwarfism resembling Meier-Gorlin syndrome. Nat Genet 43: 350–355.

Blow JJ, Gillespie PJ. 2008. Replication licensing and cancer--a fatal entanglement? Nat Rev Cancer 8: 799–806.

Burrage LC, Charng WL, Eldomery MK, Willer JR, Davis EE, Lugtenberg D, Zhu W, Leduc MS, Akdemir ZC, Azamian M et al. 2015. De Novo GMNN Mutations Cause Autosomal-Dominant Primordial Dwarfism Associated with Meier-Gorlin Syndrome. Am J Hum Genet 97: 904–913.

Cook JG. 2009. Replication licensing and the DNA damage checkpoint. Front Biosci (Landmark Ed) 14: 5013–5030.

Cook JG, Chasse DA, Nevins JR. 2004. The regulated association of Cdt1 with minichromosome maintenance proteins and Cdc6 in mammalian cells. J Biol Chem 279: 9625–9633.

Coulombe P, Gregoire D, Tsanov N, Mechali M. 2013. A spontaneous Cdt1 mutation in 129 mouse strains reveals a regulatory domain restraining replication licensing. Nat Commun 4: 2065.

Davidson IF, Li A, Blow JJ. 2006. Deregulated replication licensing causes DNA fragmentation consistent with head-to-tail fork collision. Mol Cell 24: 433–443.

De Marco V, Gillespie PJ, Li A, Karantzelis N, Christodoulou E, Klompmaker R, van Gerwen S, Fish A, Petoukhov MV, Iliou MS et al. 2009. Quaternary structure of the human Cdt1-Geminin complex regulates DNA replication licensing. Proc Natl Acad Sci U S A 106: 19807–19812.

de Munnik SA, Bicknell LS, Aftimos S, Al-Aama JY, van Bever Y, Bober MB, Clayton-Smith J, Edrees AY, Feingold M, Fryer A et al. 2012. Meier-Gorlin syndrome genotype-phenotype studies: 35 individuals with pre-replication complex gene mutations and 10 without molecular diagnosis. Eur J Hum Genet 20: 598–606.

Ferenbach A, Li A, Brito-Martins M, Blow JJ. 2005. Functional domains of the Xenopus replication licensing factor Cdt1. Nucleic Acids Res 33: 316–324.

Frigola J, He J, Kinkelin K, Pye VE, Renault L, Douglas ME, Remus D, Cherepanov P, Costa A, Diffley JFX. 2017. Cdt1 stabilizes an open MCM ring for helicase loading. Nat Commun 8: 15720.

Fujita M. 2006. Cdt1 revisited: complex and tight regulation during the cell cycle and consequences of deregulation in mammalian cells. Cell Div 1: 22.

Guernsey DL, Matsuoka M, Jiang H, Evans S, Macgillivray C, Nightingale M, Perry S, Ferguson M, LeBlanc M, Paquette J et al. 2011. Mutations in origin recognition complex gene ORC4 cause Meier-Gorlin syndrome. Nat Genet 43: 360–364.

Haland TW, Boye E, Stokke T, Grallert B, Syljuasen RG. 2015. Simultaneous measurement of passage through the restriction point and MCM loading in single cells. Nucleic Acids Res 43: e150.

Hall JR, Lee HO, Bunker BD, Dorn ES, Rogers GC, Duronio RJ, Cook JG. 2008. Cdt1 and Cdc6 are destabilized by rereplication-induced DNA damage. J Biol Chem 283: 25356–25363.

Havens CG, Walter JC. 2009. Docking of a specialized PIP Box onto chromatin-bound PCNA creates a degron for the ubiquitin ligase CRL4Cdt2. Mol Cell 35: 93–104.

Havens, CG, Walter JC. 2011. Mechanism of CRL4(Cdt2), a PCNA-dependent E3 ubiquitin ligase. Genes Dev 25: 1568–1582.

Hornbeck PV, Zhang B, Murray B, Kornhauser JM, Latham V, Skrzypek E. 2015. PhosphoSitePlus, 2014: mutations, PTMs and recalibrations. Nucleic Acids Res 43: D512–520.

Jee J, Mizuno T, Kamada K, Tochio H, Chiba Y, Yanagi K, Yasuda G, Hiroaki H, Hanaoka F, Shirakawa M. 2010. Structure and mutagenesis studies of the C-terminal region of licensing factor Cdt1 enable the identification of key residues for binding to replicative helicase Mcm proteins. J Biol Chem 285: 15931–15940.

Jin J, Arias EE, Chen J, Harper JW, Walter JC. 2006. A family of diverse Cul4-Ddb1-interacting proteins includes Cdt2, which is required for S phase destruction of the replication factor Cdt1. Mol Cell 23: 709–721.

Khayrutdinov BI, Bae WJ, Yun YM, Lee JH, Tsuyama T, Kim JJ, Hwang E, Ryu KS, Cheong HK, Cheong C et al. 2009. Structure of the Cdt1 C-terminal domain: conservation of the winged helix fold in replication licensing factors. Protein Sci 18: 2252–2264.

Lee C, Hong B, Choi JM, Kim Y, Watanabe S, Ishimi Y, Enomoto T, Tada S, Kim Y, Cho Y. 2004. Structural basis for inhibition of the replication licensing factor Cdt1 by geminin. Nature 430: 913–917.

Li C, Jin J. 2010. DNA replication licensing control and rereplication prevention. Protein Cell 1: 227–236.

Li X, Zhao Q, Liao R, Sun P, Wu X. 2003. The SCF(Skp2) ubiquitin ligase complex interacts with the human replication licensing factor Cdt1 and regulates Cdt1 degradation. J Biol Chem 278: 30854–30858.

Liontos M, Koutsami M, Sideridou M, Evangelou K, Kletsas D, Levy B, Kotsinas A, Nahum O, Zoumpourlis V, Kouloukoussa M et al. 2007. Deregulated overexpression of hCdt1 and hCdc6 promotes malignant behavior. Cancer Res 67: 10899–10909.

Liu E, Li X, Yan F, Zhao Q, Wu X. 2004. Cyclin-dependent kinases phosphorylate human Cdt1 and induce its degradation. J Biol Chem 279: 17283–17288.

Machida YJ, Teer JK, Dutta A. 2005. Acute reduction of an origin recognition complex (ORC) subunit in human cells reveals a requirement of ORC for Cdk2 activation. J Biol Chem 280: 27624–27630.

Malecki MJ, Sanchez-Irizarry C, Mitchell JL, Histen G, Xu ML, Aster JC, Blacklow SC. 2006. Leukemia-associated mutations within the NOTCH1 heterodimerization domain fall into at least two distinct mechanistic classes. Mol Cell Biol 26: 4642–4651.

Marti-Renom MA, Stuart AC, Fiser A, Sanchez R, Melo F, Sali A. 2000. Comparative protein structure modeling of genes and genomes. Annu Rev Biophys Biomol Struct 29: 291–325.

Matson JP, Dumitru R, Coryell P, Baxley RM, Chen W, Twaroski K, Webber BR, Tolar J, Bielinsky AK, Purvis JE et al. 2017. Rapid DNA replication origin licensing protects stem cell pluripotency. Elife 6.

Melixetian M, Ballabeni A, Masiero L, Gasparini P, Zamponi R, Bartek J, Lukas J, Helin K. 2004. Loss of Geminin induces rereplication in the presence of functional p53. J Cell Biol 165: 473–482.

Moreno A, Carrington JT, Albergante L, Al Mamun M, Haagensen EJ, Komseli ES, Gorgoulis VG, Newman TJ, Blow JJ. 2016. Unreplicated DNA remaining from unperturbed S phases passes through mitosis for resolution in daughter cells. Proc Natl Acad Sci U S A 113: E5757–5764.

Nevis KR, Cordeiro-Stone M, Cook JG. 2009. Origin licensing and p53 status regulate Cdk2 activity during G(1). Cell Cycle 8: 1952–1963.

Nishitani H, Sugimoto N, Roukos V, Nakanishi Y, Saijo M, Obuse C, Tsurimoto T, Nakayama KI, Nakayama K, Fujita M et al. 2006. Two E3 ubiquitin ligases, SCF-Skp2 and DDB1-Cul4, target human Cdt1 for proteolysis. EMBO J 25: 1126–1136.

Pozo PN, Cook JG. 2016. Regulation and Function of Cdt1; A Key Factor in Cell Proliferation and Genome Stability. Genes (Basel) 8.

Schindelin J, Arganda-Carreras I, Frise E, Kaynig V, Longair M, Pietzsch T, Preibisch S, Rueden C, Saalfeld S, Schmid B et al. 2012. Fiji: an open-source platform for biological-image analysis. Nat Methods 9: 676–682.

Shreeram S, Sparks A, Lane DP, Blow JJ. 2002. Cell type-specific responses of human cells to inhibition of replication licensing. Oncogene 21: 6624–6632.

Sugimoto N, Tatsumi Y, Tsurumi T, Matsukage A, Kiyono T, Nishitani H, Fujita M. 2004. Cdt1 phosphorylation by cyclin A-dependent kinases negatively regulates its function without affecting geminin binding. J Biol Chem 279: 19691–19697.

Sun J, Evrin C, Samel SA, Fernandez-Cid A, Riera A, Kawakami H, Stillman B, Speck C, Li H. 2013. Cryo-EM structure of a helicase loading intermediate containing ORC-Cdc6-Cdt1-MCM2-7 bound to DNA. Nat Struct Mol Biol 20: 944–951.

Takeda DY, Parvin JD, Dutta A. 2005. Degradation of Cdt1 during S phase is Skp2-independent and is required for efficient progression of mammalian cells through S phase. J Biol Chem 280: 23416–23423.

Truong LN, Wu X. 2011. Prevention of DNA re-replication in eukaryotic cells. J Mol Cell Biol 3: 13–22.

Vaziri C, Saxena S, Jeon Y, Lee C, Murata K, Machida Y, Wagle N, Hwang DS, Dutta A. 2003. A p53-dependent checkpoint pathway prevents rereplication. Mol Cell 11: 997–1008.

Whittaker AJ, Royzman I, Orr-Weaver TL. 2000. Drosophila double parked: a conserved, essential replication protein that colocalizes with the origin recognition complex and links DNA replication with mitosis and the down-regulation of S phase transcripts. Genes Dev 14: 1765–1776.

Yanagi K, Mizuno T, You Z, Hanaoka F. 2002. Mouse geminin inhibits not only Cdt1-MCM6 interactions but also a novel intrinsic Cdt1 DNA binding activity. J Biol Chem 277: 40871–40880.

Yekezare M, Gomez-Gonzalez B, Diffley JF. 2013. Controlling DNA replication origins in response to DNA damage - inhibit globally, activate locally. J Cell Sci 126: 1297–1306.

You Z, Ode KL, Shindo M, Takisawa H, Masai H. 2016. Characterization of conserved arginine residues on Cdt1 that affect licensing activity and interaction with Geminin or Mcm complex. Cell Cycle 15: 1213–1226.

Yuan Z, Riera A, Bai L, Sun J, Nandi S, Spanos C, Chen ZA, Barbon M, Rappsilber J, Stillman B et al. 2017. Structural basis of Mcm2-7 replicative helicase loading by ORC-Cdc6 and Cdt1. Nat Struct Mol Biol 24: 316–324.

Zhai Y, Cheng E, Wu H, Li N, Yung PY, Gao N, Tye BK. 2017. Open-ringed structure of the Cdt1-Mcm2-7 complex as a precursor of the MCM double hexamer. Nat Struct Mol Biol 24: 300–308.

